# Machine learning aided top-down proteomics on a microfluidic platform

**DOI:** 10.1101/2020.11.14.381376

**Authors:** Yuewen Zhang, Maya A. Wright, Kadi L. Saar, Pavankumar Challa, Alexey S. Morgunov, Quentin A. E. Peter, Sean Devenish, Christopher M. Dobson, Tuomas P.J. Knowles

## Abstract

The ability to determine the identity of specific proteins is a critical challenge in many areas of cellular and molecular biology, and in medical diagnostics. Here, we present a microfluidic protein characterisation strategy that within a few minutes generates a three-dimensional fingerprint of a protein sample indicative of its amino acid composition and size and, thereby, creates a unique signature for the protein. By acquiring such multidimensional fingerprints for a set of ten proteins and using machine learning approaches to classify the fingerprints, we demonstrate that this strategy allows proteins to be classified at a high accuracy, even though classification using a single dimension is not possible. Moreover, we show that the acquired fingerprints correlate with the amino acid content of the samples, which makes it is possible to identify proteins directly from their sequence without requiring any prior knowledge about the fingerprints. These findings suggest that such a multidimensional profiling strategy can lead to the development of novel method for protein identification in a microfluidic format.

## Introduction

The diverse nature of proteins and their central role in a multitude of biological processes [1–3] necessitates a requirement for highly specific and sensitive approaches for protein detection and analysis. Indeed, protein detection and characterisation approaches have been of fundamental importance for a range of biological and medical research fields and have provided valuable information for better understanding the onset of a multitude of diseases, including various forms of cancer and neurodegenerative disorders. [4–10] In particular, at the centre of the discovery of novel protein-based disease biomarkers lies the ability to identify proteins. [11–16] In this context, protein microarrays are currently one of the most widely used techniques. By providing a high spatial density array of solid-phase supported affinity reagents, such as antibodies, protein microarrays allow proteins of interest to be selectively captured and subsequently detected through the introduction of a second, frequently fluorescently labelled affinity reagent. [17,18] As such, protein microarray based approaches usually require access to two distinct antibodies each targeting a different epitope of a single protein and, similarly to any other affinity reagent mediated system, their performance is sensitive to potential undesired cross-reactivity events. On a fundamental level, such an affinity-reagent mediated strategy is inherently limited to detecting known targets for which a suitable affinity reagent was consciously included in the library and does not allow for the detection and the discovery of hitherto unknown markers.

The possibility to detect the presence of hitherto unknown targets and perform explorative screening arises when affinity-reagent free protein analysis approaches are used. In this context, various forms of mass-spectrometry have been widely used for protein identification for many decades due to their high sensitivity, resolution, accuracy and dynamic range. [19,20] In a typical experiment, fragments of proteins are formed and separated through approaches, such as liquid chromatography before their injection to a mass-spectrometer. [21–24] While top-down identification has allowed characterising a number of different protein species, its application becomes challenging in the limit of high molecular weight and low solubility species. Due to these limitations, less than 10% of mammalian proteome can be accessed through these techniques. [25] For the analysis of higher molecular weight species, bottom-up sequencing approaches have been developed, which usually involve proteolysis of a complex mixture of proteins followed by a chromatographic separation of the peptides prior to their sequencing through tandem mass spectrometry (MS/MS). Whether the analysis is performed in a top-down or bottom-up manner, mass-spectrometry generally requires extensive sample preparation steps, often resulting in significant losses, and long experimental analysis time. Moreover, the presence of less abundant species is usually masked by more abundant ones, which prevents it effective use for detecting targets that are present at low concentrations, as is the case for biomarkers during the onset and early stages of diseases. Last but not least, its operation in gas-phase, has made it challenging to extend the analysis to protein complexes that are held together through transient interactions.

Recently, approaches that would enable overcoming the drawbacks of mass spectrometry have been demonstrated and proposed. In particular, Swaminathan et al. [26] have demonstrated the possibility of immobilising peptides onto a glass slide and measuring their fluorescence through total internal reflection microscopy in consecutive cycles of Edman degradation after selectively labelling lysine and cystine residues. While demonstrating the first steps towards the feasibility of single molecule peptide fluorosequencing, [27] the approach involves a number of consecutive Edman steps, setting a limit on the speed at which the analysis can be performed.

To open up the possibility of minute-scale liquid-phase protein identification, here, we devised and demonstrated a microfluidic platform that permits the identification of protein samples on a single chip by relying on obtaining its characteristic multidimensional physicochemical signature. Specifically, by using a multi-wavelength detection system, we obtained readouts describing the tryptophan (Trp), tyrosine (Tyr) and lysine (Lys) content of the protein sample together with an estimate for their hydrodynamic radius (Figure 1). By obtaining such multidimensional signatures for a total of ten proteins and using machine learning approaches for identifying the origin of a set of validation proteins, we showed that such a strategy can be used for identifying proteins with a high confidence. The characterisation and identification process is performed on unlabelled protein samples and on a minute timescale.

**Figure 1.**
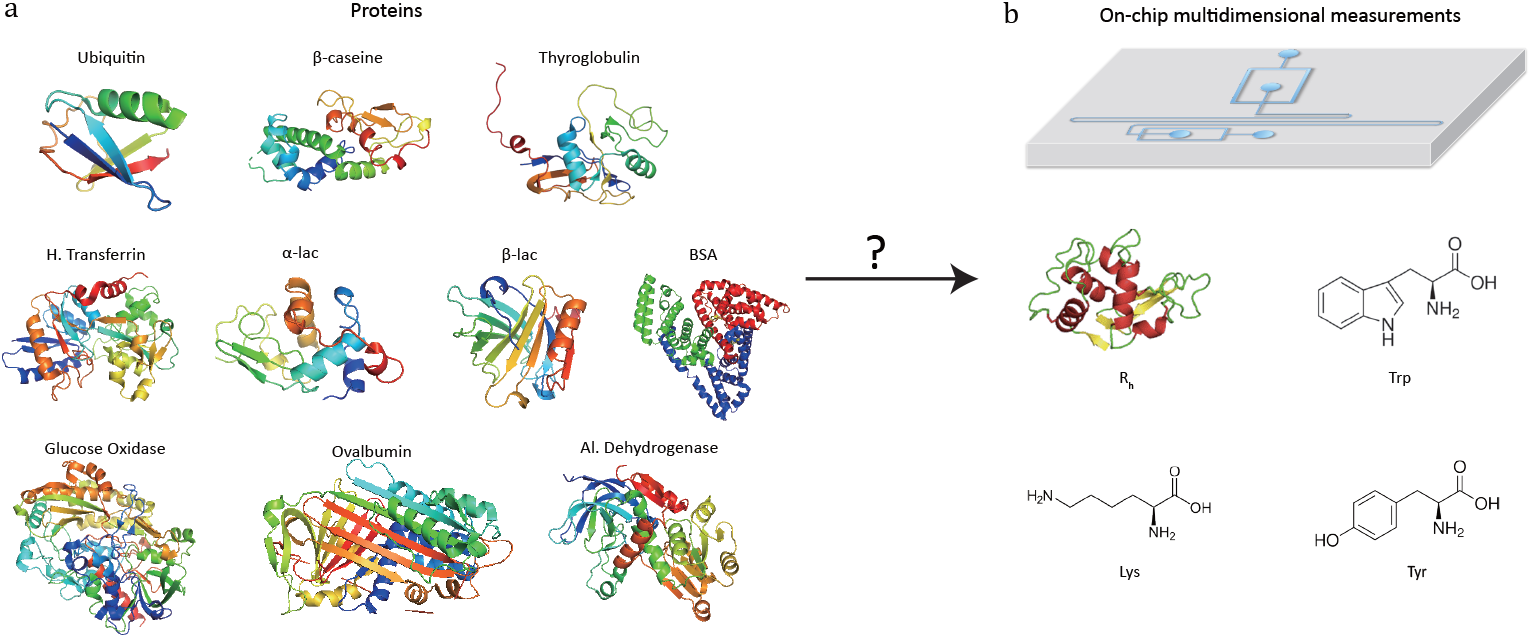
Microfluidic top down identification of proteins. **(a)** The proteins and **(b)** the microfluidic device used in this study. The device allows obtaining multi-dimensional fingerprints of protein samples that include information about their tryptophan, tyrosine and lysine content as well as the hydrodynamic radius, R_*h*_.

## Materials and Methods

### Preparation of protein samples and the labelling solution

Bovine serum albumin (BSA), *β*-lactoglobulin (b-lac), glucose oxidase, *α*-lactalbumin (a-lac), ovalbumin, human transferrin, thyroglobulin (thglb), *β*-casein and ubiquitin (ubiq) were obtained from Sigma-Aldrich, and alcohol dehydrogenase (alc, dehydr) from Alfa Aesar. All the proteins were dissolved in 25 mM phosphate buffer at pH 8.0 to a micromolar concentration range. The solution used for labelling the lysine residues (Figure 2a) included 12 mM o-phthaldialdehyde (OPA), 18 mM *β*-mercaptoethanol (BME) and 4% wt/vol sodium dodecyl sulfate (SDS) in 200 mM carbonate buffer at pH 10.5.

**Figure 2.**
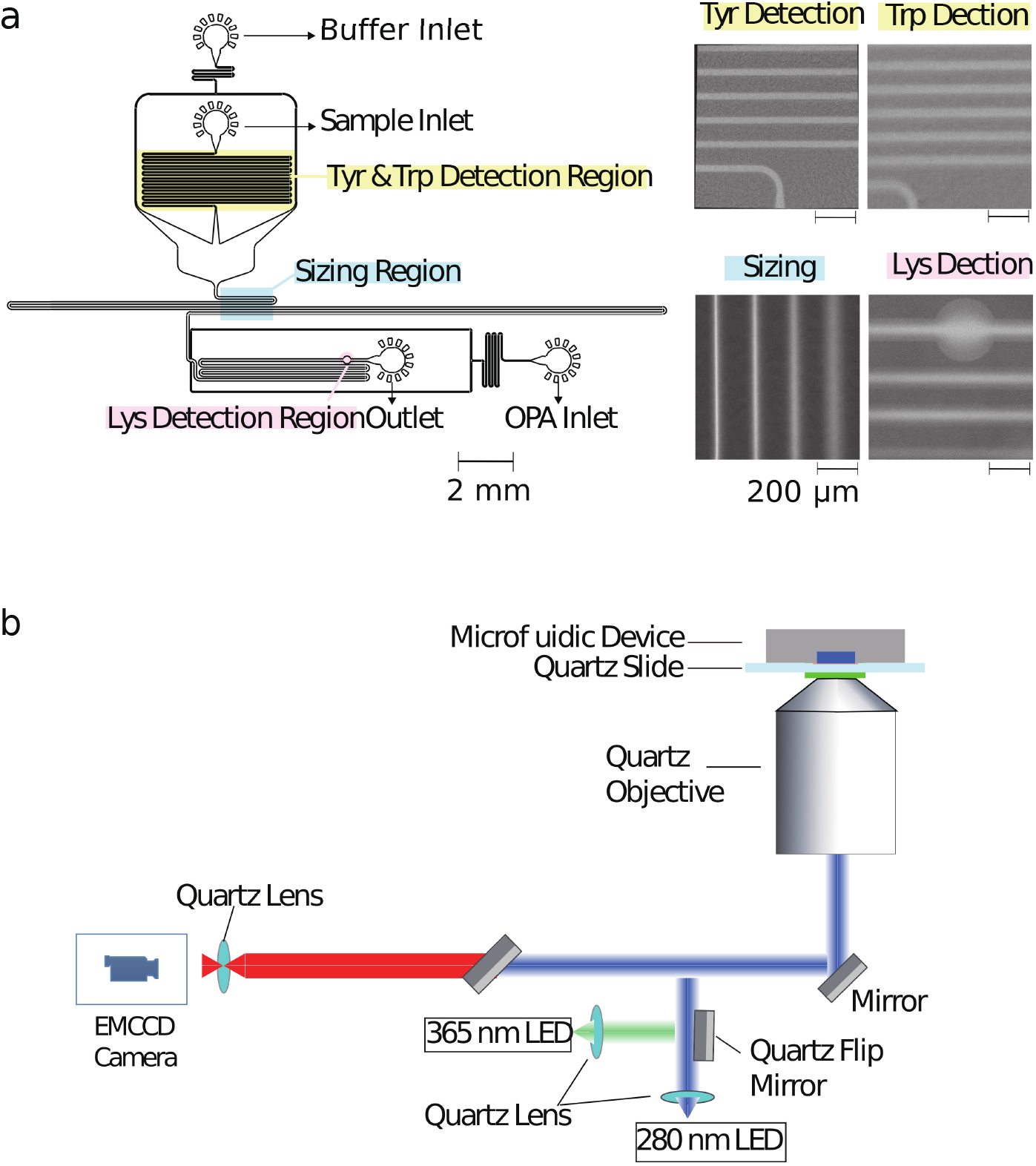
Microfluidic top-down identification strategy. **(a)** The microfluidic device used in this study allowed extracting a multidimensional characteristic signature of each analysed sample describing its (i) tryptophan (Trp) and (ii) tyrosine (Tyr) residues (yellow highlighted region), (iii) its hydrodynamic radius R_*h*_ obtained by monitoring the diffusion of the sample molecules into a co-flowing buffer (blue highlighted region) and (iv) its lysine (Lys) content (pink highlighted region). The scale bars on all insets are 200 *μ*m. **(b)** Schematic representation of the home-built inverted fluorescence microscope used. The two light sources (280 nm and 365 nm) and emission filters can be switched readily to record the characteristic fluorescent signals.

### UV-LED microscope

The schematic of the optical layout is shown in Figure 2b. The sample was excited using either a 280 nm LED (Thorlabs M280L3, UK) or a 365 nm LED (Thorlabs M365L2, UK) light source with a flip mirror used to switch between the two sources. The light from either of the LEDs was passed through an aspherical lens of focal length 20 mm to get a collimated output beam. The beam was passed through a dichroic filter cube, which consisted of an excitation filter (Semrock FF01-280/20-25) and a dichroic mirror (Semrock FF310-Di01-25×36). The light reflected by the dichroic mirror was then focussed onto the sample flowing in a microfluidic chip by an infinity corrected UV objective lens (Thorlabs LMU-10X-UVB, UK) of numerical aperture NA = 0.25. The emitted fluorescent light from the sample was collected through the same objective and an emission filter (Semrock FF01-357/44-25 for a charactersitic tryptophan, FF01-302/10-25 for a charactersitic tyrosine and FF01-452/45-25 for a charactersitic lysine signal) with an air-spaced achromatic doublet lens of focal length 20 mm (Thorlabs ACA254-200-UV) focussing it onto the camera (Rolera EMC2). All the optics used in the set-up were made out of fused silica for high transmission in the UV region. [28,29]

### Microfluidic device fabrication

The microfluidic devices were cast using polydimethylsiloxane (PDMS) (Sylgard 184 kit, Dow Corning, USA) from a silicon wafer master imprinted with 50 *μ*m high structures based using standard singlelayer soft-lithography techniques. [30] The precise height of the photoresist structures on different locations across the master mould were measured by a profilometer (DektakXT, UK) to correct the analysis for any variations in structure height across the master. Carbon black nanopowder (Sigma-Aldrich, UK) was added to the PMDS to minimise undesired autofluorescence from the PDMS devices under UV illumination during the measurements. The devices were bonded to a quartz slide (Alfa Aesar, 76.2×25.4×1.0 mm, UK) using plasma treatment (Electronic Diener Femto plasma bonder; 15 seconds at 40% of the full power). The PDMS-glass microfluidic devices were then exposed to an additional extended plasma treatment step (500 seconds at 80% of the full power) to render channel surfaces more hydrophilic with the inlets and outlets blocked with water-filled gel-loading tips immediately after the exposure to maintain their hydrophilic character.

### Device operation

To obtain a multidimensional signature for a sample, the channels of the microfluidic were first filled from the common outlet using a glass syringe (Hamilton, 500 *μ*L, UK), 27 gauge needle (Neolus Terumo, 25 gauge, 0.5 x 16 mm, UK), and polyethene tubing (Scientific Laboratory Supplies, inner diameter 0.38 mm, outer diameter 1.09 mm, UK). Gel loading tips filled with the relevant solutions were then inserted into the device inlets (Figure 2a). The fluid flow through of the solutions into the microfluidic channels was controlled using neMESYS syringe pumps (Cetoni GmbH, Germany) that was set to withdraw the solutions at a total flow rate of 200 *μ*L *h*^-1^. As described previously, [29] in order to increase the accuracy of the diffusional sizing process, the gel loading tip in the sample inlet was first filled with the auxiliary buffer and a background image of the diffusional sizing area recorded. This micrograph was later used for subtracting the static background arising from the autofluorescence of the PDMS device. The gel loading tip in the sample inlet was then carefully exchanged to a tip including the protein sample with care taken not to introduce any air bubbles in the process. For both images, an exposure time of 500 ms was used.

Finally, in order to account for any potential fluctuations in the power output of the LEDs, the intensities of standard calibration solutions (10 *μ*M L-Tryptophan and 10 *μ*M 4-methylumbelliferone both in 400 mM potassium borate buffer at pH 9.7) were recorded in a channel adjacent to the top-down identification device itself. The obtained characteristic tryptophan and tyrosine fluorescence values were then normalised by the former of this calibration readings and the lysine value by the latter of the two calibration readings.

## Results and Discussion

### Microfluidic multidimensional protein characterisation strategy

To facilitate the acquisition of multidimensional physicochemical signatures of proteins directly in solution, we designed a microfluidic device that allowed simultaneously obtaining four characteristic parameters of an unlabelled protein sample. Specifically, after introducing a sample from its corresponding inlet (Figure 2a), first, the characteristic fluorescence intensities indicative of the tryptophan and tyrosine content of the sample were recorded in the yellow highlighted area by exciting the microfluidic chip with a UV wavelength (280 nm) LED (Figure 2b) and collecting the emitted fluorescent light using two distinct filters. The filters were chosen such that the collected light originated either predominantly from its tryptophan or from its tyrosine residues (Materials and Methods).

The protein sample was then surrounded by a co-flowing buffer in order to monitor the lateral diffusion of the protein sample into an auxiliary carrier medium in space and in time. Such a strategy has been previosly shown to yield the diffusion coefficients of protein samples. [31] In particular, the device we used in this study was designed for the camera field of view (800 *μ*m × 1000 *μ*m) to include four distinct sections of this channel (blue highlighted region), so that a single image could be used to extract the diffusion coefficient as described earlier. [32] The channels were imaged using the 280 nm excitation LED in combination with the tryptophan filter as the signal from latter residue was stronger than the signal from tyrosine residues. The diffusion profiles on the micrographs were then fitted to simulated basis functions for particles of known radii and each of the simulated profiles were compared to the measured profiles in order to extract the hydrodynamic radius of each sample. [31–34]

Finally, downstream the sizing unit, an on-chip latent labelling strategy was used to conjugate the lysine residues in each protein to o-phthaldialdehyde (OPA) dye molecules [33, 35] (Materials and Methods). The characteristic fluorescence intensity from the OPA labelled lysine residues was measured (pink highlighted region) by switching the UV-LED light source to an LED light source with excitation at 365 nm wavelength (Materials and Methods) at which unconjugated OPA molecules have been observed to show only minimal background fluorescence. The dimensions of the labelling channel were chosen such that the OPA dye and the protein sample would be able to mix for over a 3 second long time period before the measurement was taken, a time scale that we had previously shown allows quantitative insight into the abundance of lysine residues in proteins. [33]

All in all, this strategy allowed us to obtain a four-dimensional signature for each protein sample using a single microfluidic chip and a dual-wavelength excitation system. One of the four parameters was later used for normalising the obtained fluorescent signals to obtain a concentration independent signature for each protein.

### Multidimensional signatures of a set of ten proteins

We analysed a set of ten different proteins (Figure 1a) and used the platform described above to obtain multidimensional signatures for each of them. In particular, we performed n = 4 repeats on all the ten proteins using a different microfluidic device for each experiment. We noted that the measured R_h_ values of all the proteins, which varied by three orders of magnitude in molecular weight, were consistent with the values reported in the literature (Supplementary Table S1).

In order to eliminate concentration dependence, the obtained signals in the tryptophan and tyrosine imaging channels were normalised by the signal in the lysine filter. This reduced the data structure to a three-dimensional signature but ensured that the obtained values were independent of the concentration of the protein that was used for analysis. Moreover, the measured intensities were corrected for fluctuations in the laser power by also measuring the fluorescence intensities of calibration solutions in a neighbouring channel, involving L-tryptophan and 4-methylumbelliferone molecules for the 280 nm and the 365 nm LED, respectively (Materials and Methods).

The characteristic spaces that each of the analysed ten proteins occupied in a three-dimensional plot are shown in Figure 3d with the 1D projections shown in Figures 3a-c and the underlying data summarised in Supplementary Table S1. In particular, the three-dimensional visualised ellipsoids (Figure 3d) were defined by the centres being the average of the four measurement points and their radii corresponding to the standard deviation of the four measurements. We noted that the ten analysed proteins varied in their physiochemical signatures with Figure 3d illustrating that it is likely that across a three-dimensional landscape each of the protein acquires a different signature.

**Figure 3.**
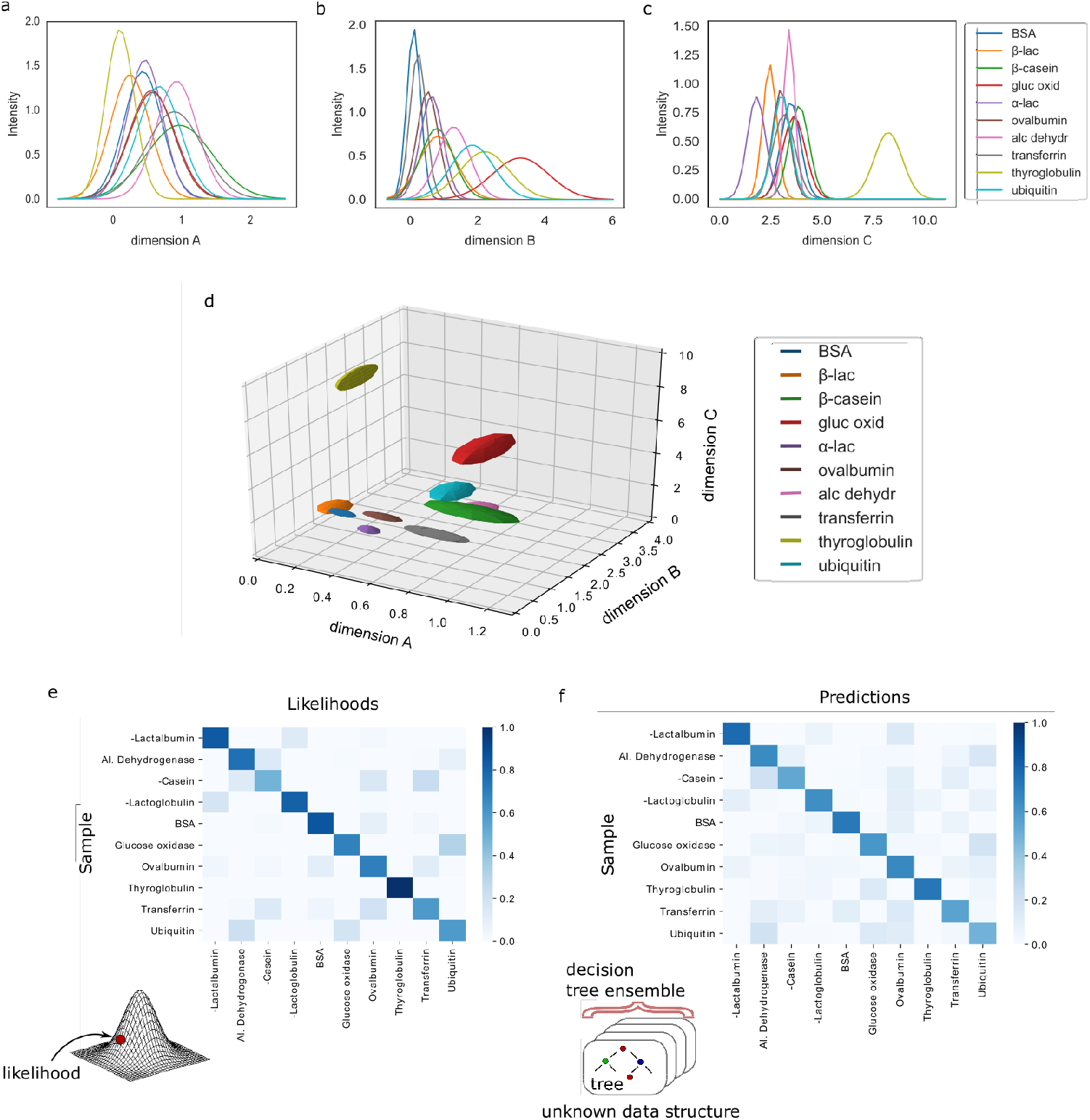
Protein classification from their multidimensional fingerprints. A set of ten proteins was profiled by acquiring their three-dimensional fingerprints described by **(a)** the ratio of the signals measured at the wavelengths where tyrosine and OPA fluoresce (Materials and Methods; dimension 1), **(b)** the ratio of the signals measured at the wavelengths where tryptophan and OPA fluoresce (dimension 2) and **(c)** the hydrodynamic radius, *R_h_* (dimension 3). All these parameters are concentration independent. **(d)** Multidimensional signatures of the proteins in a 3D space. The radii of the ellipsoids correspond to one standard deviation. **(e)** The likelihoods of protein identification and misidentification in the 3D space showed in panel **(d)** assuming multivariate Gaussian model. **(f)** The confidence levels of identification process using a random forest classifier approach that assumes no underlying data distribution. The models identified correctly 82% (multivariate Gaussian) and 90% (random forest classifier) of the tested samples.

### Protein classification

In order to evaluate whether our demonstrated platform is capable of distinguishing proteins reliably and uniquely based on their signatures, we developed two models to perform sample classification.

First, using the full data set of 10 classes of proteins with 4 experimental repeats for each class, leave-one-out cross-validation was used to assess the likelihood that a particular sample is classified as the correct protein. In particular, multivariate Gaussian distributions were fitted to each of the ten protein classes with the means computed from the four repeats within each class, or from the three remaining repeats for the class from which the validation sample was removed. The covariance matrices were computed by combining the group variance (using either four or three repeats similarly to the means) with the global variance involving the full dataset of 39 data points excluding the validation sample. A weighting factor of 0.9 was used for the group variance and a weighting factor of 0.1 for the global variance to avoid singular covariance matrices and ensure computational stability while simultaneously taking advantage of the extra information about the system as the variances in the same dimension between the different classes are likely to be similar. Finally, the likelihood of each of the validation samples belonging to each of the protein classes was calculated by estimating the probability density function of the individual multivariate Gaussians at that point.

For each protein class, the likelihood was averaged across the four experimental repeats and the resulting values were normalised to one. Figure 3e shows a heatmap of the calculated likelihoods for assigning proteins into available classes with the actual protein being measured on the vertical axis and the protein it is likely to be identified as on the horizontal axis. We observed that, individually, 33 out of 40 samples were classified correctly. Moreover, it can be seen that on average proteins are likely to be assigned to the correct class with high confidence.

We note that the above estimates were arrived at by assuming that the errors in the measurements in each dimension were normally distributed, so that the protein classes can be approximated by multivariate Gaussian distributions. In order to improve our analysis and derive an identification strategy that is not making an assumption about the distribution of the errors, we constructed a random forest classifier. As before, leave-one-out cross-validation was used on all 40 samples. In order to reduce variance, each random forest was trained with 1000 decision trees that were built using bootstrapping and with only 2 out of 3 variables selected at random to build each tree. The classification was performed using predictions by these ensemble models and, subsequently, predictions by all individual trees in the ensembles were collected to quantify the confidence of the ensemble model in making the predictions. For each group of four samples corresponding to the same protein class, the average number of trees in the ensemble predicting each target class were taken and normalised to sum to one for each protein.

Finally, a heatmap summarising the results was constructed, similarly showing the actual protein being measured on the vertical axis and the protein it is likely to be identified as on the horizontal axis (Figure 3f). The results illustrate that the model predicts the correct class of proteins with high confidence. Moreover, on the individual level, the random forest model misclassified only 4 out of 40 samples, demonstrating a superior performance to the multivariate Gaussian model. This shows that highly accurate identification of proteins is possible even when no assumptions are made about the underlying distributions of measurement errors or data structure.

### Protein identification

Having confirmed the possibility to classify an unknown protein sample by evaluating which of the pre-determined multidimensional fingerprints it resembles the most (Figure 3), we next set out to explore if it is possible to determine the origin of each of the test samples simply by performing a top-down identification process on the sample without requiring prior knowledge of the fingerprints.

To this effect, we first derived relationships that could be used to predict the sequence composition of each sample from its fingerprint. Indeed, the measured fluorescent signals were constructed in a manner where they can be expected to predominantly originate from the tyrosine, tryptophan and lysine residues of the proteins (Materials and Methods) or, in the case of the hydrodynamic radius, be linked to its molecular weight. The observed correlation between the measured fluorescent signals and the amino acid content of the proteins are shown in Figures 4a-b. As before, in order to eliminate any concentration dependence, ratios between the measured signals and amino acid compositions were used. Figure 4c additionally outlines the relationship between the measured hydrodynamic radius and the molecular weight of the proteins. We modelled all the three relationships as linear regressions with zero-intercepts and estimated the gradient of the line by minimising the ordinary least squares (Figure4a-c, dotted lines). It is possible that when more abundant data is available, more nuanced relationships between the measured signals and the sequence-specific quantities can be learned.

**Figure 4.**
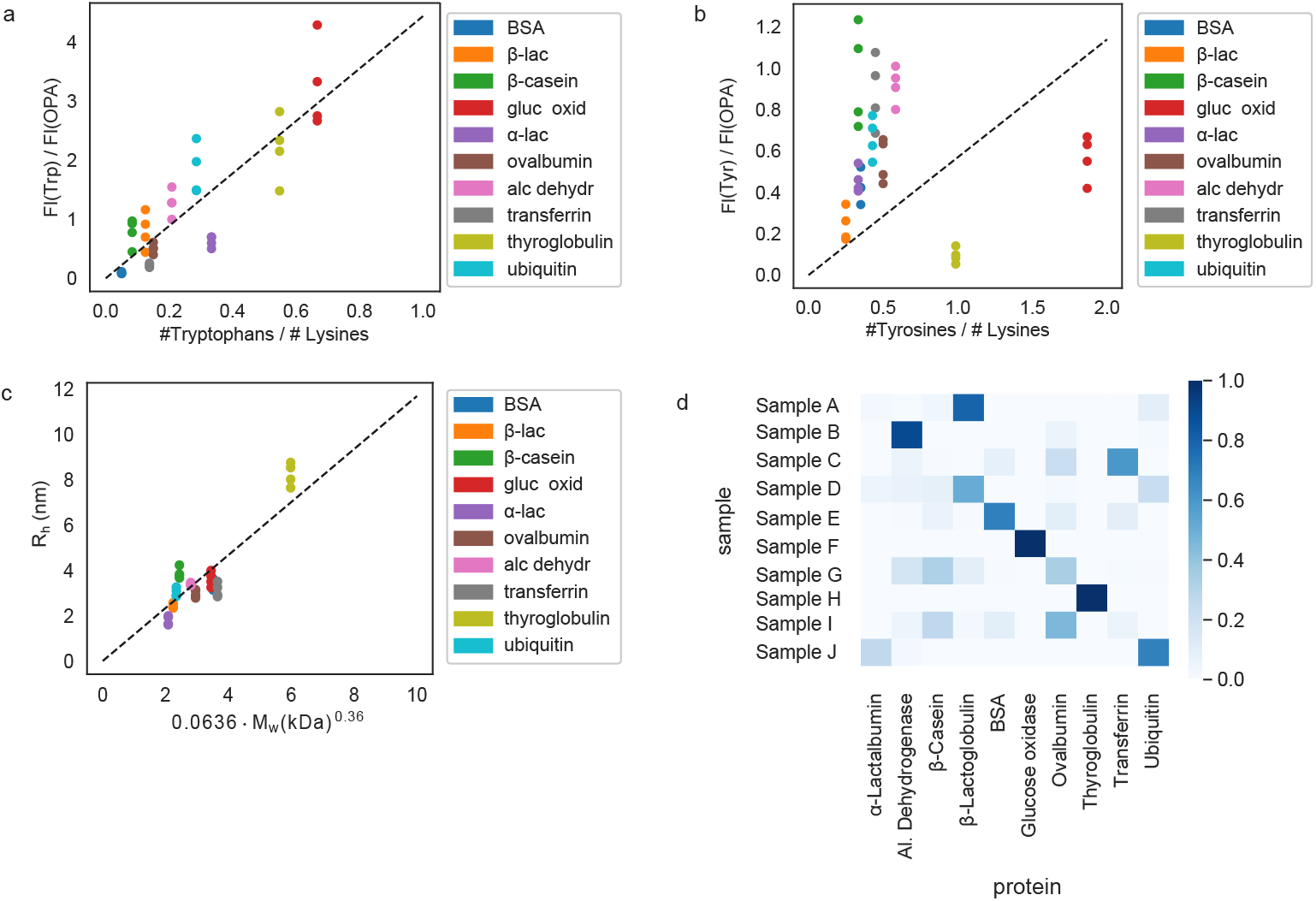
Protein identification from their sequence. The correlations between the measured signals and the sequences of the analysed proteins, specifically **(a)** the ratio of the measured tryptophan and OPA signals as a function of the tryptophan and lysine composition of the proteins, **(b)** the ratio of the measured tyrosine and OPA fluorescence signals as a function of their tyrosine and lysine composition and **(c)** the measured hydrodynamic radius, *R_h_*, as a function of the molecular weight. In all cases, the dotted line shows the best fit linear regression function with the intercept set to 0. **(d)** The measured signals for each of the ten samples (A–J) were converted to estimates of their sequence-composition using the relationships outlined in panels **(a)**-**(c)** and the latter estimates were used to evaluate the probabilities of each of the ten samples being any one of the ten proteins in our dataset by using Gaussian mixture models. The data are shown such that the correct sample appears on the diagonal of the matrix. Individual samples were identified correctly on 21 out of 40 occasions. When averaging the results over *n* = 4 repeats, seven out of ten proteins were identified correctly.

Next, the derived relationships (Figure 4a-c) were used to convert the measured three-dimensional signature of our test samples into their predicted 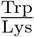 and 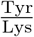 ratios and molecular weights. To eliminate any information leakage, we re-fitted the linear regression by excluding the test sample. Following this step, the z-score of the measured sample being a particular protein in the database was calculated by using the estimated sequence-specific properties of this sample as the mean value and the measurement noise as the standard deviation. The heatmap describing the most likely origin of each of the test samples is shown in Figure 4d with the data arranged such that the correct sample appeared on the diagonal of the matrix. These data show that, on average, seven out the ten proteins were identified correctly. On the level of individual measurements, samples were identified correctly in 21 out of 40 experiments (Figure 5). These results illustrate that not only can the multidimensional signatures used for classifying proteins into pre-determined clusters (Figure 3e-f), it is also possible to convert the measured signals into absolute sequence-specific parameters and through this process identify the test samples.

**Figure 5.**
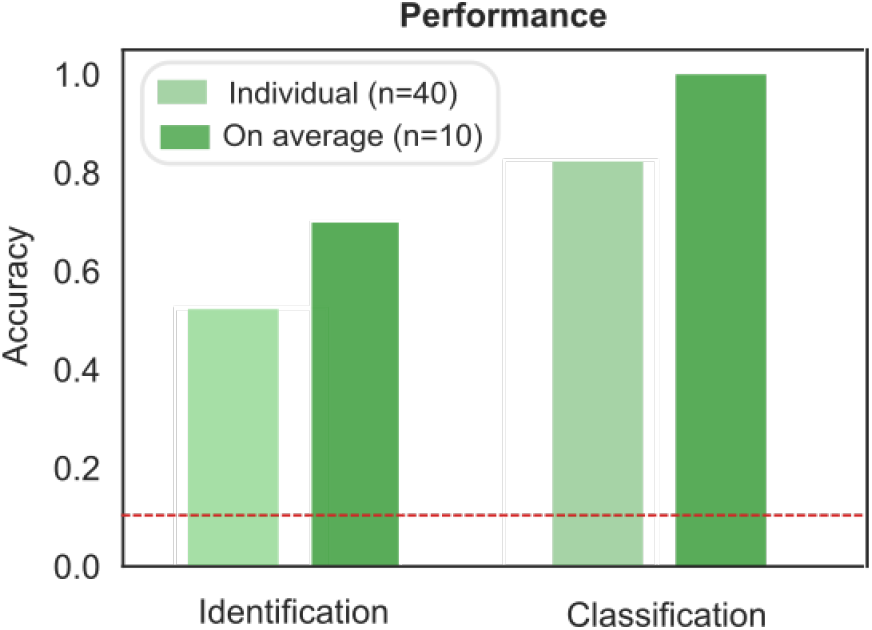
Comparison of the performance of the protein classification (Figure 3) and identification (Figure 4) strategies. When identifying a measured sample directly from its sequence, samples were identified correctly on 53% of the occasions or on 70% of the occasions when the results were averaged across the four repeats performed on each samples. When pre-determined fingerprints were used, proteins were classified correctly on 83% of the occasions or on 100% of the occasion when the results were averaged across the repeats. The red dotted line corresponds to the case where the classification or identification was performed by a process of random guessing.

## Conclusions

We developed a strategy for obtaining multidimensional physicochemical signatures of individual proteins on a single microfluidic chip and showed that this strategy can be used for protein classification as well as top-down identification. Specifically, we achieved this objective by designing a chip on which the hydrodynamic radius of a protein sample could be obtained simultaneously with signals describing its tryptophan, tyrosine and lysine content. We showed that this approach generated unique fingerprints for all the ten proteins in our test set and, moreover, that the signatures can be used to identify proteins in a top-down manner. Our results suggest that an on-chip multidimensional protein characterisation strategy could serve as a powerful probe-free approach for on-chip biomarker profiling using only microlitre sized samples.

## Author Contributions

Y.Z., M.A.W., K.L.S., P.C. and T.P.J.K. designed the study and conceived the experiments, P.C. built the experimental hardware, Y.Z. and M.A.W. acquired the data and processed the micrographs for which Q.A.E.P. contributed relevant software. K.L.S. and A.S.M. analysed and interpreted the data and built the machine learning models. Y.Z., K.L.S., A.S.M. and T.P.J.K. wrote the original manuscript and M.A.W., P.C., S.D., Q.A.E.P. and C.M.D. reviewed it.

## Acknowledgements

This is a contribution from the Cambridge Centre for Misfolding Diseases. The research leading to these results has received funding from the ERC under the European Union Seventh Framework Programme (FP7/2007-2013) through the ERC grant PhysProt(agreement n° 337969’ T.P.J.K.), the EPSRC (K.L.S), the Schmidt Science Fellows program, in partnership with the Rhodes Trust (K.L.S), the BBSRC (C.M.D, T.P.J.K), and the Frances and Augustus Newman Foundation (T.P.J.K.).

